# Development and implementation of an integrated preclinical atherosclerosis database

**DOI:** 10.1101/2023.09.12.557423

**Authors:** Rachel Xiang, Yihua Wang, Megan M. Shuey, Brigett Carvajal, Quinn S. Wells, Joshua A. Beckman, Iris Z. Jaffe

## Abstract

**Background:** Basic scientists have used preclinical animal models to explore mechanisms driving human diseases for decades, resulting in thousands of publications, each supporting causative inferences. Despite substantial advances in the mechanistic construct of disease, there has been limited translation from individual studies to advances in clinical care. An integrated approach to these individual studies has the potential to improve translational success.

**Methods:** Using atherosclerosis as a test case, we extracted data from the two most common mouse models of atherosclerosis (ApoE and LDLR knockout). We restricted analyses to manuscripts published in two well-established journals, *Arteriosclerosis, Thrombosis, and Vascular Biology* and *Circulation*, as of query in 2021. Predefined variables including experimental conditions, intervention and outcomes were extracted from each publication to produce a preclinical atherosclerosis database.

**Results:** Extracted data include animal sex, diet, intervention type and distinct plaque pathologies (size, inflammation, lipid content). Procedures are provided to standardize data extraction, attribute interventions to specific genes and transform the database for use with available transcriptomics software. The database integrates hundreds of genes, each directly tested in vivo for causation in a murine atherosclerosis model. The database is provided to allow the research community to perform integrated analyses that reflect the global impact of decades of atherosclerosis investigation.

**Conclusions:** Future database uses include interrogation of sub-datasets associated with distinct plaque pathologies, cell-type or sex. We provide the methods and software needed to apply this approach to the extensive repository of peer-reviewed data utilizing preclinical models to interrogate mechanisms of diverse human diseases.

## Introduction

For decades, basic scientists have used preclinical animal models to explore mechanisms that may drive human diseases. The most commonly used preclinical species is the murine model chosen for rapid breeding, small size, relatively low cost, and ease of genetic manipulation. As examples, there are 26 mouse heart failure models,^1^ 11 abdominal aortic aneurysm models,^2^ as well as non-cardiovascular disease mouse models to study Alzheimer’s Disease (12 models),^3^ lung cancer (41 models)^4^ among many others.

Atherosclerosis, the vascular disease that causes myocardial infarction and stroke in humans, is the leading cause of death worldwide. The two most common mouse models of atherosclerosis are the apolipoprotein E (ApoE) and LDL receptor (LDLR) knockout (KO) models.^5^ These mice develop atherosclerotic plaques with pathology that mimics many aspects of the human disease, particularly when fed an atherogenic high fat diet. Since the development of these models in the 1990s, over 10,000 manuscripts have been published.^6^ Each manuscript typically examines the impact of at least one individual perturbation on atherosclerosis parameters in the setting of KO of either ApoE or LDLR, providing new insights into the molecular mechanisms driving atherogenesis. Unlike human genetic analyses of genetic variants and atherosclerosis phenotypes that can only show association, pharmacological and genetic KO studies in mouse models compared to placebo or gene-intact controls, facilitates the evaluation of causal relationships between specific interventions with atherosclerotic phenotypes. However, this reductionist approach is not without limitations. Genetic KO models in mice, while specific in targeting one gene, may perturb entire pathways leading to the ultimate phenotype observed. Additionally, sequential studies that perturb individual genes may be subject to redundancy, as multiple tested targets may converge on a common disease mechanism. Moreover, gene deletion does not faithfully recapitulate complex human genetics. Indeed, few individual targets identified from preclinical mouse investigations have translated into novel therapies for human disease.^5^

The growing body of preclinical literature combined with the rapid advance of systems biology methods presents the possibility of synthesizing large amounts of preclinical information to identify biological pathways and master regulators which may have greater potential as therapeutic targets. Such approaches would immediately add value to the existing preclinical data, acquired over many decades at considerable cost, by providing a synthesis of causal relationships associated with atherosclerosis pathogenesis. The lack of a standardized method to aggregate preclinical data for systems biology approaches prevents the full utilization of available peer-reviewed investigator-generated data and represents an unmet gap in extracting the full insight from this work. We recently developed a database structure and data extraction method to integrate preclinical data for pathway and network analysis using atherosclerosis as a test case. The novelty of this database construction and method of integration lies in the potential for repurposing decades of preclinical investigations, each of which describe a causal relationship between a single-gene perturbation and atherosclerotic plaque endpoints. This approach has potential to uncover novel insights into mechanisms driving atherosclerotic disease through integrated analysis and to enhance the translational relevance of available preclinical investigations. Our described database structure and methods may also be extended to any preclinical disease model for which large amounts of data have been published.

## Methods and Results

### PubMed Query

To identify published atherosclerosis studies using the ApoE knockout (KO) and LDLR-KO mouse models, the EndNote and PubMed databases were queried using the search strategy summarized in the **Table**. Results from this search strategy in EndNote and PubMed were combined, with duplicates removed, resulting in identification of more than 6000 manuscripts from the ApoE query and over 4000 manuscripts from the LDLR query (numbers change on weekly basis as new papers are published). For the proof-of-concept study, we extracted all data published from 1995 (date of first publication of ApoE-KO mouse) through 2020 in the journals *Arteriosclerosis, Thrombosis, and Vascular Biology (ATVB)* and *Circulation*, a total of 1535 manuscripts (**Figure 1A**).

**Table 1:**
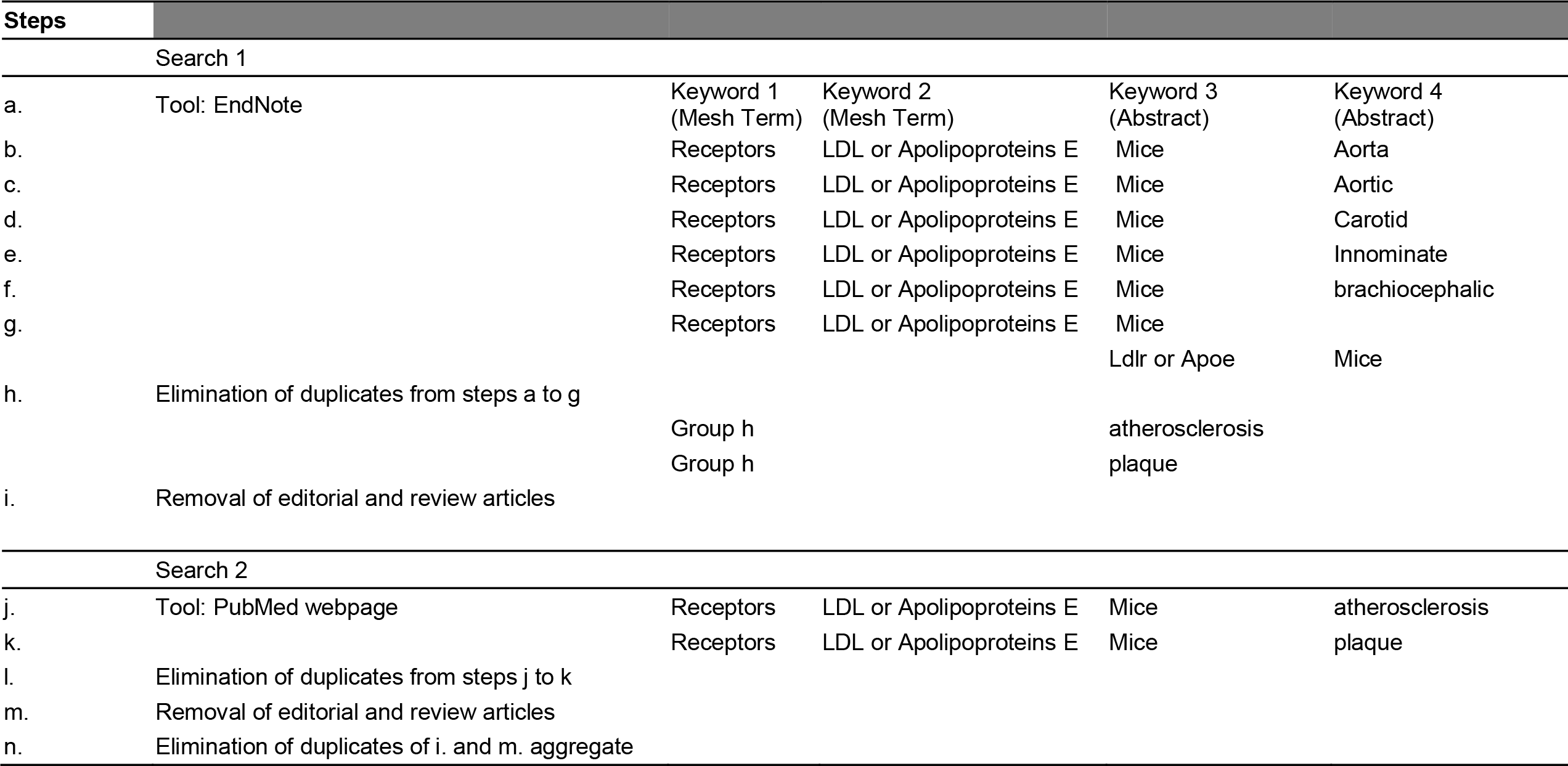
Search strategy within selected journals for models of choice.

**Figure 1.**
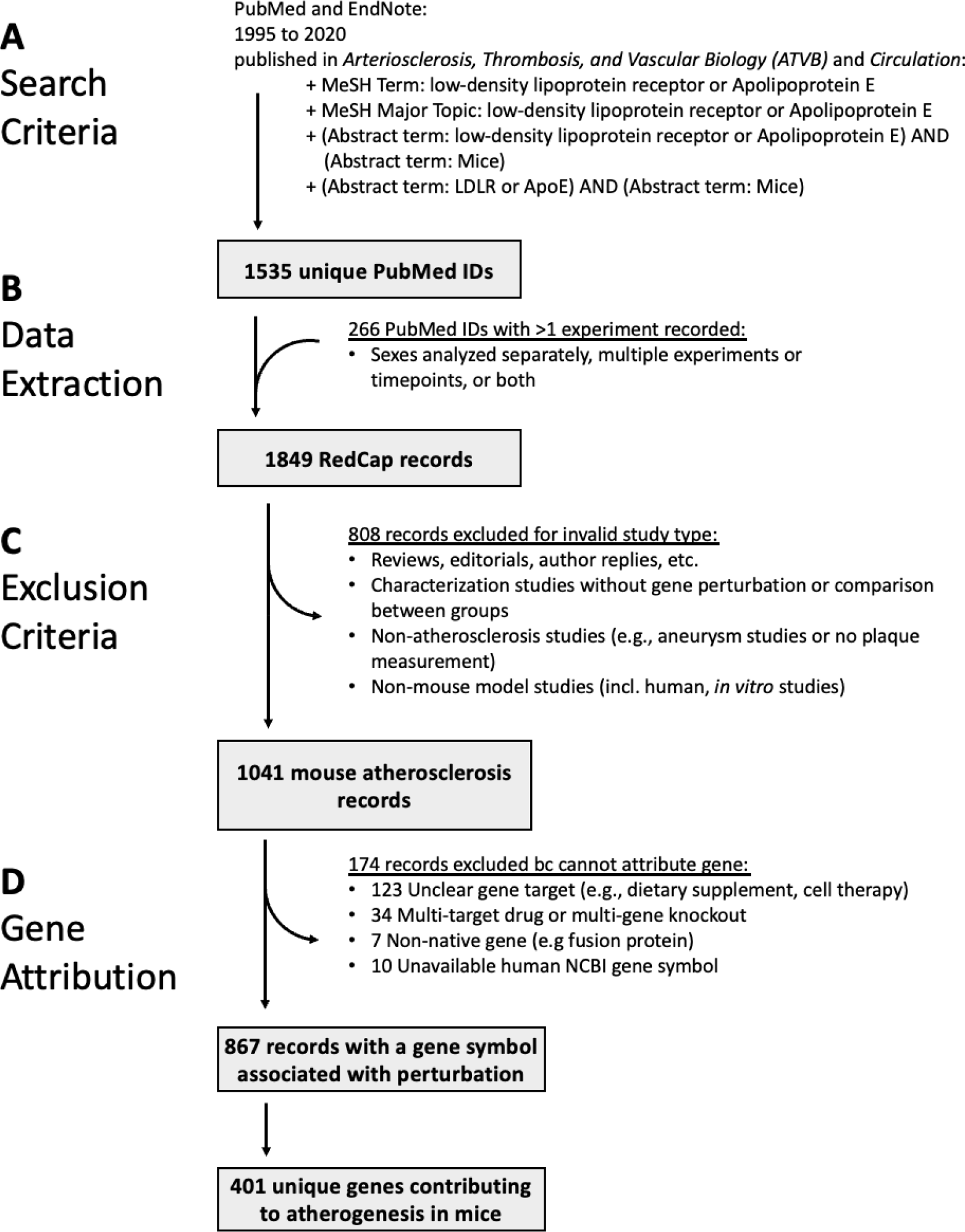
Flow diagram of database development from manuscript identification to identification of gene attribution. The first step of the process is **A)** the identification of manuscripts from two journals using predefined search criteria. Next, **B)** data is extracted from the 1535 manuscripts to identify specific experimental designs, interventions, and outcomes. Manuscripts that included more than one experimental design and outcome pairing were uploaded as separate RedCap records that were specific to the specific experimental variables. From the 1849 records, **C)** 808 (43.7%) were excluded due to identification as an invalid study type, including: a review, editorial, non-murine study, or other criteria. Finally, in **D)** the intervention strategies are attributed to specific genes, example: results corresponding to administration of an angiotensin converting enzyme (ACE) inhibitor would be attributed to the *Ace* gene.

These journals were chosen for their rigorous peer-review process, large volumes of papers published in the disease models of interest, and consistent publication of atherosclerosis studies over the full time frame. This search method may be adapted to the research interest or disease model of choice.

### Establishing Variables and Developing the Database Architecture

The Research Electronic Data Capture (REDCap) web application was used to build and manage data extraction as it allows data entry by multiple users over time with the ability to export a single integrated preclinical database. For manuscript identification the PubMed identification (PMID) number, manuscript title, and year of publication were recorded. To collect experimental design variables, the REDCap form utilized drop down menus and text fields to collect the key data components including: mouse atherosclerosis model (ApoE-KO or LDLR-KO), sex of the animals (male, female, both, not indicated), duration of the study (number of weeks), and type of diet fed to the mice (normal chow, high fat diet). Details regarding the experimental intervention used in each study were collected including: the mode of perturbation (i.e., drug, siRNA, viral transduction, genetic KO, cell-specific genetic KO), drug dosage, cell type of the genetic KO (or whole body), NCBI gene symbol of the gene/protein impacted, and whether the impact of the intervention on gene function was gain of function (i.e., transgenic overexpression, activator drug) or loss of function (gene knockout, siRNA knock down, inhibitor drug). Finally, data fields were included to collect information regarding the impact of the experimental perturbation on atherosclerosis phenotypes. This included the location where atherosclerotic plaque measurements were made (aorta and/or aortic branch vessel (carotid, innominate, brachiocephalic)). Additionally, three atherosclerosis plaque phenotypes were captured: atherosclerotic plaque size, plaque inflammation, and plaque lipid content. Since the magnitude of changes in these parameters are highly influenced by differences in experimental design (e.g., drug dose, experiment duration, diet) and quantification methods (e.g., plaque burden along the whole aorta verses plaque size in the aortic root or brachiocephalic artery), the impact of each perturbation on each plaque variable was standardized by recording in the form of the direction of the effect on atherosclerosis (increase = +1, decrease = -1, no change =0), rather than the magnitude. In cases where multiple results using different experimental conditions were reported within a single manuscript (e.g., each sex reported separately, two time points measured, a gene KO and a drug), each set of outcomes was recorded as an independent REDCap record (**Figure 1B**). To distinguish between multiple entries from the same manuscript, a suffix was appended to the respective PMID (i.e., M, F for animal sex, #1, 2, 3 for different experimental conditions).

### Building an Integrated Mouse Preclinical Atherosclerosis Database

Data were manually extracted from each manuscript into one or more REDCap forms (**Figure 1**). The search criteria identified 1,535 manuscripts (**Figure 1A, Supplemental Table 1**) from which 17% included multiple experiments resulting in a total of 1,849 total REDCap records (**Figure 1B**).

#### 1. Extractor Training and Quality Control

Research staff were trained to follow an established protocol to extract targeted information from each manuscript. Training required completion of a training set consisting of 10-20 manuscripts from which all variables were previously extracted into a master database and used as the gold standard to verify accuracy. Training set results were compared to the master and any discrepancies clarified with each trainee. The training was repeated with additional sets of manuscripts until >98% concordance was achieved. This training process was employed to achieve interrater reliability and consistency during data extraction. Going forward after training, 5% of manuscripts from the total continued to be extracted by two investigators to address drift and further confirm and maintain consistency over time.

#### 2. Inclusion/Exclusion Criteria

Records were excluded from further analysis if no perturbation was performed (e.g., reviews, editorials, characterization) or if no atherosclerosis phenotype was measured (see **Figure 1C** for specific exclusion criteria). The preclinical atherosclerosis database comprised a total of 1,041 REDCap records after application of inclusion and exclusion criteria.

#### 3. Database Export

The finalized REDCap database was exported in .csv format and is available for download (see **Supplemental Database**). Within this database, 97% of the records indicated an impact on plaque size, 63% on inflammation, and 24% on plaque lipid content (**Figure 2A**). Many studies measured multiple plaque phenotypes, with the most common being plaque size and inflammation (44%) and an additional 16% measuring all three phenotypes.

**Figure 2.**
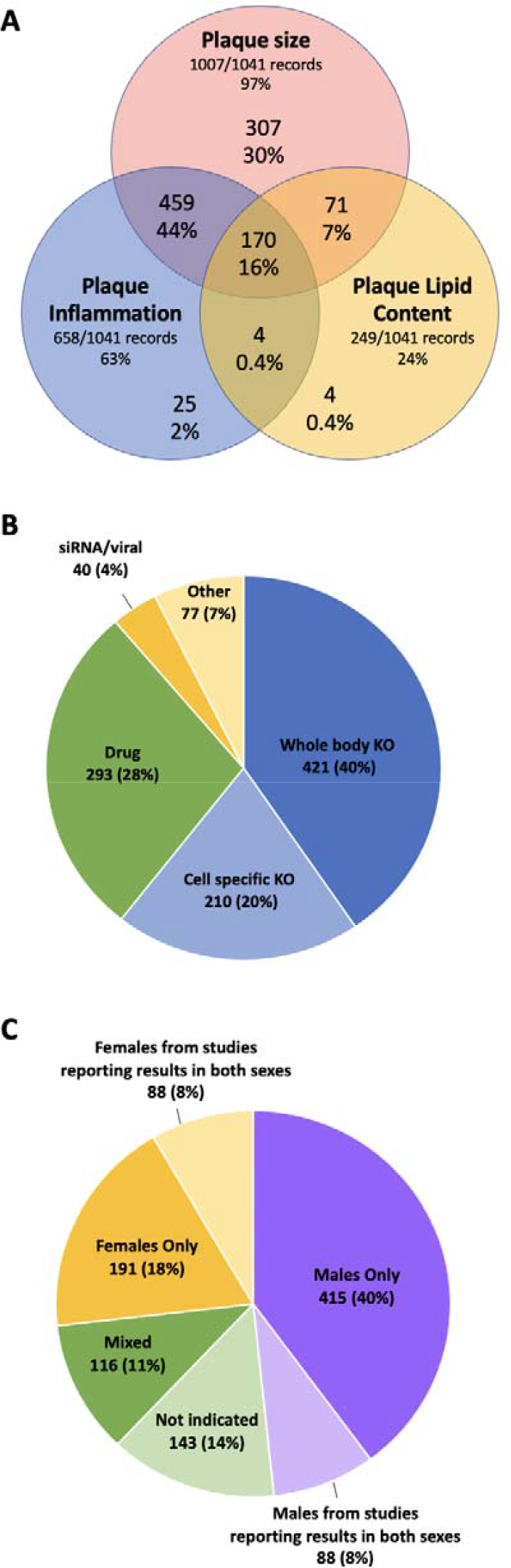
Breakdown of database descriptors based on specific outcomes, intervention type, and sex-specific experimental design. **A)** Demonstrates the number of records in the database that studied plaque size, inflammation, or lipid content as an outcome, as well as those that studied multiple outcomes, e.g. 170 (16%) studied all three. **B)** The number of studies that utilized a specific intervention type including: whole body or cell-specific KO, drug, siRNA/viral, or other. **C)** The number of studies in the database that used only one murine model sex to complete experiments as well as those for whom sex was not indicated (143 studies (14%).

### Post-export Data Processing for Integrated Analysis

A broad array of software are available, either freely or with a subscription, to investigate gene ontology, pathways, networks, and biological functions that are statistically enriched in “omics” datasets. Most were designed to input transcriptomic data in which changes in gene expression between ≥2 conditions are available across many genes. The preclinical atherosclerosis dataset was further processed to generate a file format that can be uploaded into such analysis software as described below.

#### 1. Gene attribution

Standardized methods were developed to attribute a specific target gene to each perturbation in each published murine atherosclerosis study (**Figure 1D**). For genetic KO models (60% of all records, **Figure 2B**) and viral/siRNA knock down or overexpression studies (4% of all records), gene attribution was self-evident, and the human NCBI gene symbol for the targeted gene was attributed as causative. For drug studies, the gene symbol for the intended drug target was used (e.g,, if an angiotensin converting enzyme (ACE) inhibitor was given to mice, then the ACE gene was attributed). This paradigm was used despite the known limitation that drugs are not completely specific, particularly at high doses. No NCBI gene was included in the pathway analysis for 174 records (17% of records in the database) where there was no clear gene target (i.e., dietary supplement, multi-targeted drug, deletion of a cell type, expression of a non-native or fusion protein). The final dataset yielded 867 records, including 401 unique genes, in which a single gene perturbation was identified that was associated with a change in at least one plaque parameter in the ApoE-KO or LDLR-KO mouse model.

#### 2. Data transformation for pathway analysis software

The original REDCap dataset describes the impact of a gain or loss of function of a gene on plaque measurement endpoints. To prepare our dataset for common pathway analysis software compatibility, the data were converted into a change in gene expression that would associate with a positive impact on each atherosclerosis measurement. To do this, all atherosclerosis endpoint measurements were converted to the positive direction, and values describing the direction of gene regulation were converted accordingly. For example, if a drug KO (loss-of-function) resulted in less inflamed plaques, then an increase in plaque inflammation was expected with increased expression of the gene. These transformations resulted in a dataset assigning each gene symbol to its predicted direction of regulation given an increase for each plaque measurement (plaque size, inflammation, lipid content). In this way, the transformed dataset was analogous to the typical large gene expression datasets required for pathway analysis software. An R script was developed to automatically convert the REDCap exported file into a database that is compatible with pathway analysis software, automating the post-export processing and transformation steps. The R script is available in the **Supplemental Information** of this paper. This transformation is not disease specific and can be applied to other databases generated by this method.

### Integrated Pre-clinical Atherosclerosis Dataset Description and Analysis Opportunities

#### 1. Atherosclerosis measurements (**Figure 2A**)

Virtually all of the studies included in the database (97%, 1,007/1,041), provided data on the impact of a perturbation on atherosclerotic plaque size or burden. This provides a dataset sufficiently large for pathway and network analysis to identify upstream regulators or global processes and functions that may drive plaque development. In humans, atherosclerotic plaques are generally asymptomatic until hemodynamically significant or plaque erosion or rupture leads to acute thrombotic events, i.e., myocardial infarction and ischemic stroke. Human pathology studies show that more inflamed plaques with higher lipid content are more vulnerable to rupture, leading to adverse cardiovascular events.^7^ From our dataset, 658 (63%) of records reported data on inflammation, allowing for distinct analyses to understand what may drive inflammation in atherosclerosis, a process which may be more likely to link to human outcomes data. Once a plaque ruptures, the pro-thrombogenic, lipid laden core is exposed to blood and induces thrombosis. The dataset includes only 249 records (24%) in which lipid content was measured, extraction of additional manuscripts to produce a larger dataset is likely needed to interrogate specific regulators of plaque lipid content.

#### 2. Modes of perturbation (Figure 2B)

Interrogation of the database reveals that 60% of records come from genetic knockout studies (40% whole body KO and 20% cell-specific KO). An additional 4% used siRNA or viral transduction to knock down or overexpress a gene of interest, with drug administration (28%) and other perturbations (7%) making up the rest of the database. Genetic KO studies offer the greatest degree of precision and confidence in gene attribution, as most murine KO studies provide molecular confirmation of the specificity of gene deletion. Hence, a distinct analysis using only data from KO studies may limit false attribution, and could be compared with the complete dataset to determine whether this might have greater fidelity for translation. Cell-type specific KO studies comprise 20% of the data collected, with increasing frequency over time, as floxed mice and specific Cre recombinase driver mice have become more commonly available. These include deletions specific to myeloid cells, macrophages, T-cells, endothelial cells, smooth muscle, liver, adipose and others. As the available data accumulates using these models, the potential to perform cell-type specific pathway and network analysis for atherosclerosis drivers will become a possibility.

#### 3. Animal Sex (Figure 2C)

Despite known sex differences in the prevalence and impact of atherosclerosis in males versus females, our dataset reveals that sex-differences are still rarely examined. Overall, 40% of the records provide results in only male animals and 18% in only females. An addition 11% of studies mix data from the two sexes and 14% do not indicate the sex of the animals used. Hence over half of studies include only one sex and almost a quarter studies provide no information about which sex may be impacted by the perturbation studied. Only 8% of the studies provided data in both sexes. Those studies yielded 16% of the records, as the results were extracted as separate records for each sex, with some showing concordance and some discordance of the outcome by sex. This finding could help explain some limitations in translation as findings identified in only one sex in mice are tested in human trials that mix participants of different genders. In addition to providing insights about research practices with regards to sex as a biological variable, larger datasets of this type would allow for sex-specific analyses to identify sex-specific pathways or networks that might nominate precision medicine strategies leading to potential for sex-specific therapeutic trials.

## Discussion

In summary, we have developed a method to extract and integrate results from hundreds to thousands of published manuscripts to generate an integrated preclinical database (**Figure 3**). Using atherosclerosis as a test case, we extracted data from the two most common mouse models of atherosclerosis (ApoE and LDLR KO models) to produce a preclinical atherosclerosis database that integrates hundreds of genes, each of which has been directly tested in vivo for causation in a murine atherosclerosis model. The database provides substantial opportunities for integrated analysis to identify pathways, networks, and upstream regulators that reflect the integrated impact of decades of data. Using this database, we have already identified novel pathways and networks associated with atherogenesis.^8^ By combining the integrated preclinical results with clinical databases that link genetically predicted gene expression to human atherosclerosis phenotypes, we recently demonstrated that pathway level analysis yields greater correlation of genes with human atherosclerosis phenotypes than individual genes tested in mice.^8^ We have also used this method to compare different preclinical models and found that data extracted from ApoE-KO and LDLR-KO mouse models converge on similar pathways, despite testing distinct genes, with no model showing superiority. Future uses of the dataset include interrogation of sub-datasets associated with distinct plaque pathologies, (inflammation, lipid content), cell-type specific analysis, and sex-specific findings. Finally, we provide the methods and code needed to apply this approach to the extensive repository of peer reviewed data already published utilizing preclinical models to interrogate mechanisms of diverse human diseases.

**Figure 3.**
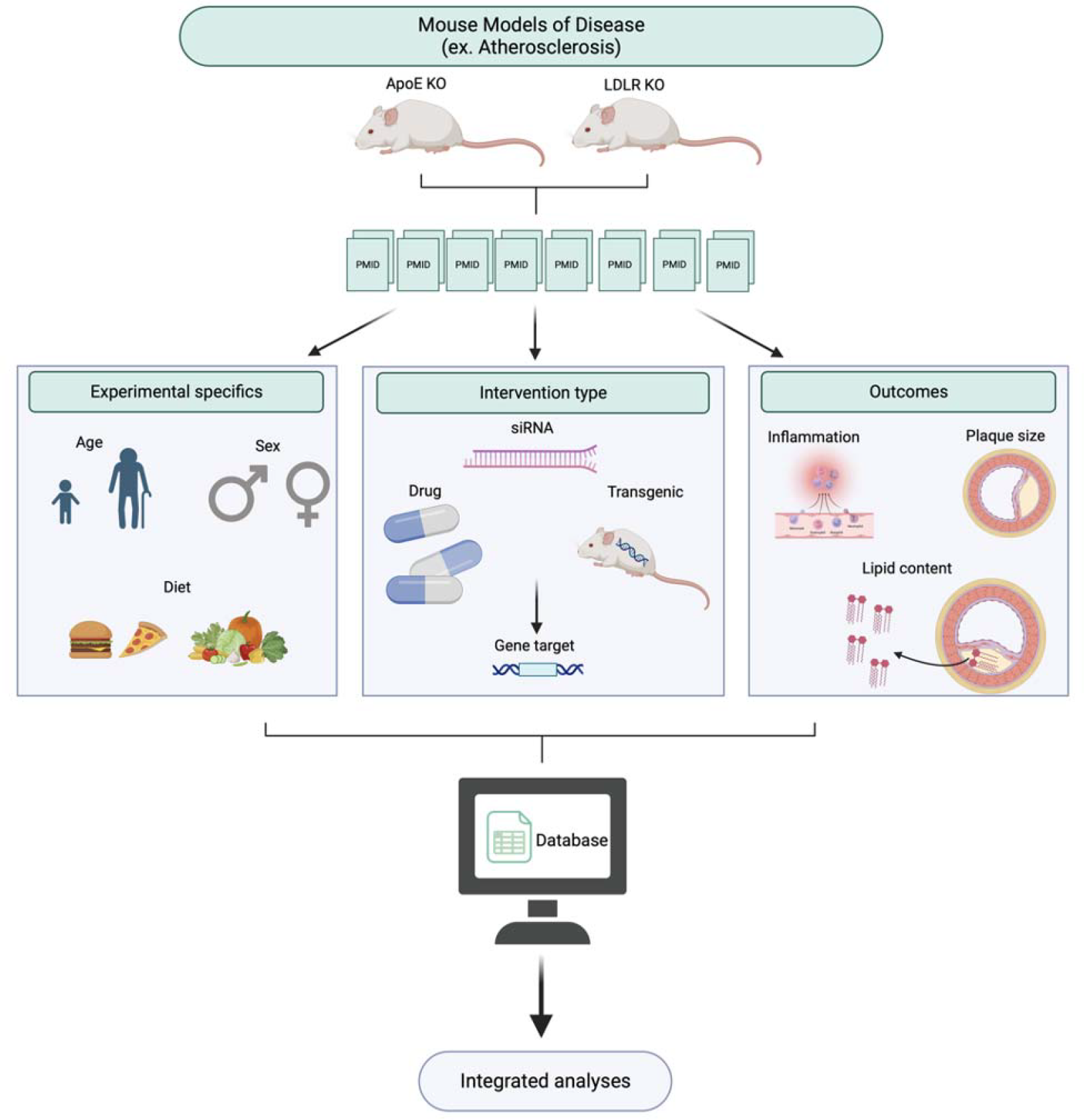
Diagram of the process of database collection, variable extraction, and development with the goal of using the outputs for integrated analyses. We demonstrate the step-wise approach for the development of a database for mouse models for disease with the goals of these variables being used for future integrated analyses such as pathway and network discovery. This diagram demonstrates the approach used in our example for two murine models of atherosclerosis (ApoE and LDLR), however, the same logic and variable

National Institutes of Health (NIH) support has resulted in the creation large amounts of biomedical research data including quantitative and qualitative datasets from fundamental research using model organisms, clinical, observational and epidemiological studies. In June of 2018, NIH released its Strategic Plan for Data Science. NIH defines data science as “the interdisciplinary field of inquiry in which quantitative and analytical approaches, processes, and systems are developed and used to extract knowledge and insights from increasingly large and/or complex sets of data.”^9^ To accomplish its strategic plan, the NIH created the Big Data to Knowledge program to maximize and accelerate the development of innovative and transformative approaches for extant big data. Our work extends this opportunity to also include the tens of thousands of fundamental preclinical model investigations already completed, peer reviewed, and published by organizing that data into a dataset amenable to use with data science approaches.

Mouse models of atherosclerosis, particularly genetic knockouts, have provided a robust platform for interrogation of the mechanisms of disease. More than 10,000 manuscripts where either apolipoprotein E (ApoE) or the low-density lipoprotein receptor (LDLR) have been knocked out (KO) have been published.^10-12^ The benefit of these mice is the routine development of atherosclerotic plaque that resemble those in humans, particularly when fed a western or high-fat diet.^13^ These mouse platforms have permitted the testing of dietary, genetic, environmental, and pharmacological perturbations in a stable genetic background. Moreover, there is evidence that coronary artery disease pathways derived from human genome-wide association studies (GWAS) show a strong overlap with mouse GWAS for atherosclerosis.^14^ Leveraging these models, thousands of studies have implicated individual interventions as directly modulating development, progression, and severity of atherosclerosis in mice.

The potency of a single genetic knockout facilitates disease mechanism evaluation but has some notable limitations. By design, mouse investigations interrogate single targets at specific time points, with a specific diet, and, commonly, with only a single sex investigated. Using ApoE and LDLR KO models, the exploration of varying diets, drug interventions and gene perturbations on atherosclerosis plaque size, inflammation, and lipid content have yielded a series of insights that have advanced the understanding of atherogenesis, but primarily through singular advances variably incorporated by the community into an inchoate aggregate. For complex, multigenic disorders that involve large numbers of genes, current single-study approaches have not facilitated a smooth integration into a model network of contributing genetic factors. As a result, the relevance of animal models for common human disorders has been questioned, largely based on the modest track record of drug targets developed from animal models that show efficacy in humans.^15-18^ The Preclinical Science Integration and Translation (PRESCIANT) method coalesces all investigations from a series of single observations to consideration of the totality of work.^8^

The creation of this novel dataset builds on the mission outlined by the NIH. First, we seek to maximize the extraction of information from data already created, vetted by reviewers, and published in scientific journals. This upcycling demonstrates that the value of the work in advancing discovery need not end soon after publication. Second, although we focused on atherosclerosis, the fundamental methods of our database creation may be applied to any disease with preclinical modeling. The report of our search terms, the depositing of our dataset, and the instructions for generating files amenable to assessment of distilled data using available data analysis software and pipelines, provide a roadmap for use by others. Finally, our published work^8^ shows one method by which these data can be translated to humans, using human databases linking genetics to clinical phenotypes. We anticipate the use of data like these may spark other uses to further enhance the information recovery from completed work.

## Conclusion

The extension of “big data” methods to fundamental, basic science investigation provides an additional avenue of exploration, discovery, and capitalization of completed scientific experimentation. Our method of extracting and integrating published preclinical data highlights the value of each individual study through holistic interrogation of the body of work. We believe that this tool may help to bridge the current gap between preclinical model exploration and human disease and treatment.

## Supporting information

Supplemental Database

## Acknowledgments

None

