## Supplemental Methods for "Development and implementation of an integrated preclinical atherosclerosis database"

R script for redcap database management and variable extraction

##should manually check redcap output. Running this script assumes that redcap export is formatted and accurate.

##this performs basic cleanup and will pull some demographics data

##more nuanced demographics will still have to be extracted by hand.

\## ----load libraries, set wd---------------------------------------------------

library(magrittr)

library(tidyverse)

library(dplyr)

library(readr)

library(readxl)

library(ggplot2)

library(ggVennDiagram)

setwd("~/Documents/presciant") #set working directory to folder containing redcap export

\## ----upload RedCap output as .csv---------------------------------------------

redcap_df <- read.csv("ApoE_DATA_2022-03-09_1237.csv")

cat("Total number of records:", nrow(redcap_df))

\## ----basic cleanup functions, generate and save athero dataset as .csv -------

redcap_df[redcap_df == ''] <- NA

redcap_df[redcap_df == 'N/A'] <- NA

redcap_df[redcap_df == 'ND'] <- NA

athero_only <- redcap_df

athero_only <- athero_only %>%

filter_at(vars(delta_lesion_size,delta_plaque_inflammation,delta_lipid_oro_content_of), any_vars(!is.na(.)))

view(athero_only)

cat("# records with recorded athero measurements:", nrow(athero_only))

write.csv(athero_only, file = "redcap_athero_measaured.csv")

\## ----get demographics for athero only dataset---------------------------------

##basic animal sex:

cat("# female records:", nrow(filter(athero_only, animal_sex == 'FEMALE')))

cat("# male records:", nrow(filter(athero_only, animal_sex == 'MALE')))

cat("# mixed records:", nrow(filter(athero_only, animal_sex == "BOTH")))

cat("# not indicated records:", nrow(filter(athero_only, is.na(animal_sex))) + nrow(filter(athero_only, animal_sex == 'NOT INDICATED')))

##model:

#apoe

cat("# ApoE athero records:", nrow(filter(ew_athero_only, atherosclerosis_model == 'APOE KO'| atherosclerosis_model == "ApoE KO")))

apoe_athero_only <- athero_only

apoe_athero_only <- apoe_athero_only %>%

filter(atherosclerosis_model == "APOE KO")

write.csv(apoe_athero_only, file = "apoe_athero_records.csv")

#ldlr

cat("# LDLR athero records:", nrow(filter(ew_athero_only, atherosclerosis_model == 'LDLR KO')))

ldlr_athero_only <- athero_only

ldlr_athero_only <- ldlr_athero_only %>%

filter(atherosclerosis_model == "LDLR KO")

write.csv(ldlr_athero_only, file = "ldlr_athero_records.csv")

#other mouse athero model

cat("# other mouse model records:", nrow(filter(athero_only, atherosclerosis_model == 'OTHER MOUSE ATHERO MODEL')))

##intervention type

#total

cat("# drug:", nrow(filter(athero_only, study_type == 'DRUG')))

cat("# full body knockout:", nrow(filter(athero_only, study_type == 'KO')))

cat("# cell specific knockout:", nrow(filter(athero_only, study_type == 'Cell specific KO')))

cat("# siRNA/viral:", nrow(filter(athero_only, study_type == 'siRNA/viral')))

cat("# other:", nrow(filter(athero_only, is.na(study_type))))

#by model (i.e. apoe and ldlr separately)

#location athero measured

cat("# aorta:", nrow(filter(athero_only, location_athero_quantified == 'AORTA')))

cat("# aortic branch:", nrow(filter(athero_only, location_athero_quantified == 'AORTIC BRANCH')))

cat("# other / not indicated:", nrow(filter(athero_only, is.na(location_athero_quantified))))

#plaque measurements - returns crude venn diagram of plaque measurements

has_plaque_measurements <- athero_only[, c("delta_lesion_size", "delta_plaque_inflammation", "delta_lipid_oro_content_of")]

has_plaque_measurements[!is.na(has_plaque_measurements)] <- 1

view(has_plaque_measurements)

ggVennDiagram(lapply(has_plaque_measurements, function(x) which(x==1)))

#high fat diet related **all valid entries should be in WEEKS. probably best to do manually

hfd_length <- athero_only[, c("weeks_high_fat_diet")]

hfd_length <- sapply(hfd_length, as.numeric)

hist(hfd_length)

\## ----transformation for IPA---------------------------------------------------

\# in athero df, if gene_symbol is valid (not blank, not ND), add dataframe column to numerize GOF/LOF as 1/-1 --> multiply plaque measurements by this (add 3 cols)

\# all of the gene names need in NCBI format otherwise will fail & no multiple genes per entry

df_single_gene <- athero_only %>%

filter_at(vars(gene_symbol), any_vars(!is.na(.)))

df_single_gene <- df_single_gene %>%

filter_at(vars(impact_of_intervention_on), any_vars(!is.na(.)))

view(df_single_gene)

df_single_gene$delta_lesion_size[is.na(df_single_gene$delta_lesion_size)] <- 999

df_single_gene$delta_plaque_inflammation[is.na(df_single_gene$delta_plaque_inflammation)] <- 999

df_single_gene$delta_lipid_oro_content_of[is.na(df_single_gene$delta_lipid_oro_content_of)] <- 999

df_single_gene$delta_lesion_size <-as.numeric(df_single_gene$delta_lesion_size)

df_single_gene$delta_plaque_inflammation <-as.numeric(df_single_gene$delta_plaque_inflammation)

df_single_gene$delta_lipid_oro_content_of <-as.numeric(df_single_gene$delta_lipid_oro_content_of)

df_single_gene$gene_FC <- with(df_single_gene, ifelse(impact_of_intervention_on == 'GOF', 1, -1))

df_single_gene$geneFC_size <- df_single_gene$delta_lesion_size * df_single_gene$gene_FC

df_single_gene$geneFC_inflam <- df_single_gene$delta_plaque_inflammation * df_single_gene$gene_FC

df_single_gene$geneFC_lipid <- df_single_gene$delta_lipid_oro_content_of * df_single_gene$gene_FC

view(df_single_gene)

df_ipa <- df_single_gene %>% select(pmid, gene_symbol, geneFC_size, geneFC_inflam, geneFC_lipid)

df_ipa[df_ipa == 999] <- NA

df_ipa[df_ipa == -999] <- NA

#check and write as .csv

view(df_ipa)

write.csv(df_ipa, file = "data_ipa_upload.csv")
