## Supplemental Table for "Development and implementation of an integrated preclinical atherosclerosis database"

### Slide 1
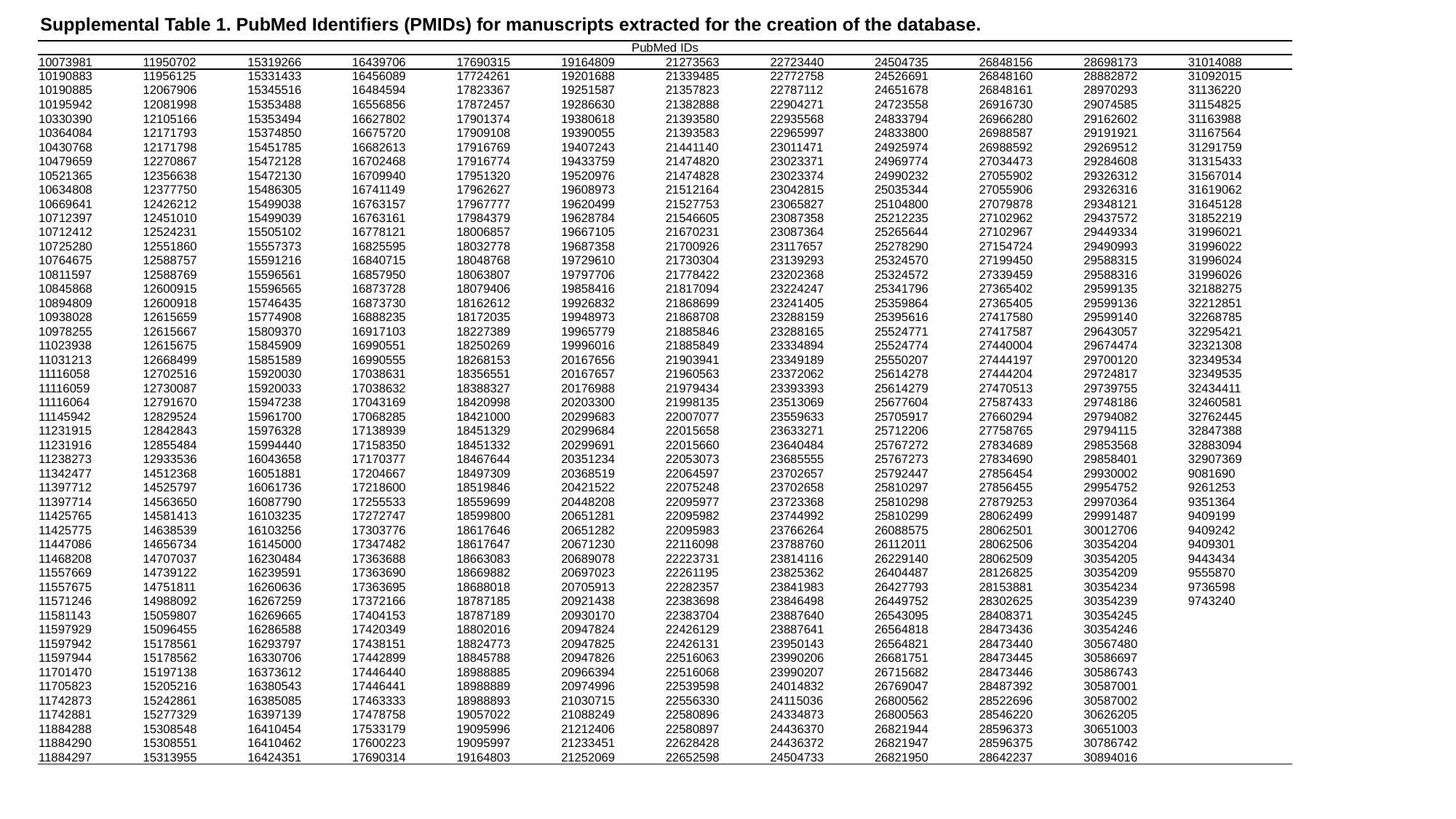

Supplemental Table 1. PubMed Identifiers (PMIDs) for manuscripts extracted for the creation of the database.
| PubMed IDs | | | | | | | | | | | |
| --- | --- | --- | --- | --- | --- | --- | --- | --- | --- | --- | --- |
| 10073981 | 11950702 | 15319266 | 16439706 | 17690315 | 19164809 | 21273563 | 22723440 | 24504735 | 26848156 | 28698173 | 31014088 |
| 10190883 | 11956125 | 15331433 | 16456089 | 17724261 | 19201688 | 21339485 | 22772758 | 24526691 | 26848160 | 28882872 | 31092015 |
| 10190885 | 12067906 | 15345516 | 16484594 | 17823367 | 19251587 | 21357823 | 22787112 | 24651678 | 26848161 | 28970293 | 31136220 |
| 10195942 | 12081998 | 15353488 | 16556856 | 17872457 | 19286630 | 21382888 | 22904271 | 24723558 | 26916730 | 29074585 | 31154825 |
| 10330390 | 12105166 | 15353494 | 16627802 | 17901374 | 19380618 | 21393580 | 22935568 | 24833794 | 26966280 | 29162602 | 31163988 |
| 10364084 | 12171793 | 15374850 | 16675720 | 17909108 | 19390055 | 21393583 | 22965997 | 24833800 | 26988587 | 29191921 | 31167564 |
| 10430768 | 12171798 | 15451785 | 16682613 | 17916769 | 19407243 | 21441140 | 23011471 | 24925974 | 26988592 | 29269512 | 31291759 |
| 10479659 | 12270867 | 15472128 | 16702468 | 17916774 | 19433759 | 21474820 | 23023371 | 24969774 | 27034473 | 29284608 | 31315433 |
| 10521365 | 12356638 | 15472130 | 16709940 | 17951320 | 19520976 | 21474828 | 23023374 | 24990232 | 27055902 | 29326312 | 31567014 |
| 10634808 | 12377750 | 15486305 | 16741149 | 17962627 | 19608973 | 21512164 | 23042815 | 25035344 | 27055906 | 29326316 | 31619062 |
| 10669641 | 12426212 | 15499038 | 16763157 | 17967777 | 19620499 | 21527753 | 23065827 | 25104800 | 27079878 | 29348121 | 31645128 |
| 10712397 | 12451010 | 15499039 | 16763161 | 17984379 | 19628784 | 21546605 | 23087358 | 25212235 | 27102962 | 29437572 | 31852219 |
| 10712412 | 12524231 | 15505102 | 16778121 | 18006857 | 19667105 | 21670231 | 23087364 | 25265644 | 27102967 | 29449334 | 31996021 |
| 10725280 | 12551860 | 15557373 | 16825595 | 18032778 | 19687358 | 21700926 | 23117657 | 25278290 | 27154724 | 29490993 | 31996022 |
| 10764675 | 12588757 | 15591216 | 16840715 | 18048768 | 19729610 | 21730304 | 23139293 | 25324570 | 27199450 | 29588315 | 31996024 |
| 10811597 | 12588769 | 15596561 | 16857950 | 18063807 | 19797706 | 21778422 | 23202368 | 25324572 | 27339459 | 29588316 | 31996026 |
| 10845868 | 12600915 | 15596565 | 16873728 | 18079406 | 19858416 | 21817094 | 23224247 | 25341796 | 27365402 | 29599135 | 32188275 |
| 10894809 | 12600918 | 15746435 | 16873730 | 18162612 | 19926832 | 21868699 | 23241405 | 25359864 | 27365405 | 29599136 | 32212851 |
| 10938028 | 12615659 | 15774908 | 16888235 | 18172035 | 19948973 | 21868708 | 23288159 | 25395616 | 27417580 | 29599140 | 32268785 |
| 10978255 | 12615667 | 15809370 | 16917103 | 18227389 | 19965779 | 21885846 | 23288165 | 25524771 | 27417587 | 29643057 | 32295421 |
| 11023938 | 12615675 | 15845909 | 16990551 | 18250269 | 19996016 | 21885849 | 23334894 | 25524774 | 27440004 | 29674474 | 32321308 |
| 11031213 | 12668499 | 15851589 | 16990555 | 18268153 | 20167656 | 21903941 | 23349189 | 25550207 | 27444197 | 29700120 | 32349534 |
| 11116058 | 12702516 | 15920030 | 17038631 | 18356551 | 20167657 | 21960563 | 23372062 | 25614278 | 27444204 | 29724817 | 32349535 |
| 11116059 | 12730087 | 15920033 | 17038632 | 18388327 | 20176988 | 21979434 | 23393393 | 25614279 | 27470513 | 29739755 | 32434411 |
| 11116064 | 12791670 | 15947238 | 17043169 | 18420998 | 20203300 | 21998135 | 23513069 | 25677604 | 27587433 | 29748186 | 32460581 |
| 11145942 | 12829524 | 15961700 | 17068285 | 18421000 | 20299683 | 22007077 | 23559633 | 25705917 | 27660294 | 29794082 | 32762445 |
| 11231915 | 12842843 | 15976328 | 17138939 | 18451329 | 20299684 | 22015658 | 23633271 | 25712206 | 27758765 | 29794115 | 32847388 |
| 11231916 | 12855484 | 15994440 | 17158350 | 18451332 | 20299691 | 22015660 | 23640484 | 25767272 | 27834689 | 29853568 | 32883094 |
| 11238273 | 12933536 | 16043658 | 17170377 | 18467644 | 20351234 | 22053073 | 23685555 | 25767273 | 27834690 | 29858401 | 32907369 |
| 11342477 | 14512368 | 16051881 | 17204667 | 18497309 | 20368519 | 22064597 | 23702657 | 25792447 | 27856454 | 29930002 | 9081690 |
| 11397712 | 14525797 | 16061736 | 17218600 | 18519846 | 20421522 | 22075248 | 23702658 | 25810297 | 27856455 | 29954752 | 9261253 |
| 11397714 | 14563650 | 16087790 | 17255533 | 18559699 | 20448208 | 22095977 | 23723368 | 25810298 | 27879253 | 29970364 | 9351364 |
| 11425765 | 14581413 | 16103235 | 17272747 | 18599800 | 20651281 | 22095982 | 23744992 | 25810299 | 28062499 | 29991487 | 9409199 |
| 11425775 | 14638539 | 16103256 | 17303776 | 18617646 | 20651282 | 22095983 | 23766264 | 26088575 | 28062501 | 30012706 | 9409242 |
| 11447086 | 14656734 | 16145000 | 17347482 | 18617647 | 20671230 | 22116098 | 23788760 | 26112011 | 28062506 | 30354204 | 9409301 |
| 11468208 | 14707037 | 16230484 | 17363688 | 18663083 | 20689078 | 22223731 | 23814116 | 26229140 | 28062509 | 30354205 | 9443434 |
| 11557669 | 14739122 | 16239591 | 17363690 | 18669882 | 20697023 | 22261195 | 23825362 | 26404487 | 28126825 | 30354209 | 9555870 |
| 11557675 | 14751811 | 16260636 | 17363695 | 18688018 | 20705913 | 22282357 | 23841983 | 26427793 | 28153881 | 30354234 | 9736598 |
| 11571246 | 14988092 | 16267259 | 17372166 | 18787185 | 20921438 | 22383698 | 23846498 | 26449752 | 28302625 | 30354239 | 9743240 |
| 11581143 | 15059807 | 16269665 | 17404153 | 18787189 | 20930170 | 22383704 | 23887640 | 26543095 | 28408371 | 30354245 | |
| 11597929 | 15096455 | 16286588 | 17420349 | 18802016 | 20947824 | 22426129 | 23887641 | 26564818 | 28473436 | 30354246 | |
| 11597942 | 15178561 | 16293797 | 17438151 | 18824773 | 20947825 | 22426131 | 23950143 | 26564821 | 28473440 | 30567480 | |
| 11597944 | 15178562 | 16330706 | 17442899 | 18845788 | 20947826 | 22516063 | 23990206 | 26681751 | 28473445 | 30586697 | |
| 11701470 | 15197138 | 16373612 | 17446440 | 18988885 | 20966394 | 22516068 | 23990207 | 26715682 | 28473446 | 30586743 | |
| 11705823 | 15205216 | 16380543 | 17446441 | 18988889 | 20974996 | 22539598 | 24014832 | 26769047 | 28487392 | 30587001 | |
| 11742873 | 15242861 | 16385085 | 17463333 | 18988893 | 21030715 | 22556330 | 24115036 | 26800562 | 28522696 | 30587002 | |
| 11742881 | 15277329 | 16397139 | 17478758 | 19057022 | 21088249 | 22580896 | 24334873 | 26800563 | 28546220 | 30626205 | |
| 11884288 | 15308548 | 16410454 | 17533179 | 19095996 | 21212406 | 22580897 | 24436370 | 26821944 | 28596373 | 30651003 | |
| 11884290 | 15308551 | 16410462 | 17600223 | 19095997 | 21233451 | 22628428 | 24436372 | 26821947 | 28596375 | 30786742 | |
| 11884297 | 15313955 | 16424351 | 17690314 | 19164803 | 21252069 | 22652598 | 24504733 | 26821950 | 28642237 | 30894016 | |
